# Haystack: systematic analysis of the variation of epigenetic states and cell-type specific regulatory elements

**DOI:** 10.1101/199067

**Authors:** Luca Pinello, Rick Farouni, Guo-Cheng Yuan

**Affiliations:** Department of Molecular Pathology, Massachusetts General Hospital, Boston MA, USA; Harvard Medical School, Boston MA, USA; Dana-Farber Cancer Institute, Boston MA, USA; Harvard School of Public Health, Boston MA, USA

## Abstract

**Motivation:** With the increasing amount of genomic and epigenomic data in the public domain, a pressing challenge is how to integrate these data to investigate the role of epigenetic mechanisms in regulating gene expression and maintenance of cell-identity. To this end, we have implemented a computational pipeline to systematically study epigenetic variability and uncover regulatory DNA sequences that play a role in gene regulation.

**Results:** Haystack is a bioinformatics pipeline to characterize hotspots of epigenetic variability across different cell-types as well as cell-type specific cis-regulatory elements along with their corresponding transcription factors. Our approach is generally applicable to any epigenetic mark and provides an important tool to investigate cell-type identity and the mechanisms underlying epigenetic switches during development. Additionally, we make available a set of precomputed tracks for a number of epigenetic marks across several cell types. These precomputed results may be used as an independent resource for functional annotation of the human genome.

**Availability:** The Haystack pipeline is implemented as an open-source, multiplatform, Python package called haystack_bio available at <u>https://github.com/pinellolab/haystack_bio</u>.

## 1 Introduction

Epigenetic patterns are highly cell type specific, and influence gene expression programs (Jenuwein and Allis, 2001). Recently, a large amount of epigenomic data across many cell types has been generated and deposited in the public domain, in part thanks to large consortia such as Roadmap Epigenomics Project (Bernstein, *et al.*., 2010), and ENCODE (Dunham, *et al.*, 2012). These data sources offer unprecedented opportunities for systematic integration and comparison. In an earlier work (Pinello, *et al.*, 2014), we developed and validated a computational strategy to systematically evaluate cross-cell-type epigenetic variability and to identify the underlying regulatory factors of such variability. Here we provide an implementation of this strategy that automatically integrates multiple data types in an easy to use command line software. Our goal is to facilitate biologists’ efforts at analyzing epigenetic data without the burden of coding, and to enable researchers to integrate their own sequencing data with information from the public domain.

## 2 Description

Haystack takes as input the genome-wide distributions of an epigenetic mark across multiple cell types or subjects measured by ChIP-seq, DNase-seq, ATAC-seq or similar assays as well as gene expression profiles quantified by microarray or RNA-seq. Users can start with publicly available preprocessed data or integrate their own data in the pipeline by providing BAM or bigWig files that can be generated by existing tools such as the ENCODE Uniform Processing Pipelines. Haystack’s entire computational pipeline can be executed with a single command (i.e. *haystack_pipeline*). The pipeline is composed of three modules: *haystack_hotspots*, *haystack_motifs*, and *haystack_tf_activity_plane*. Each module is designed to carry out a distinct but related task (Fig. 1A), as described below.

**Fig. 1.**
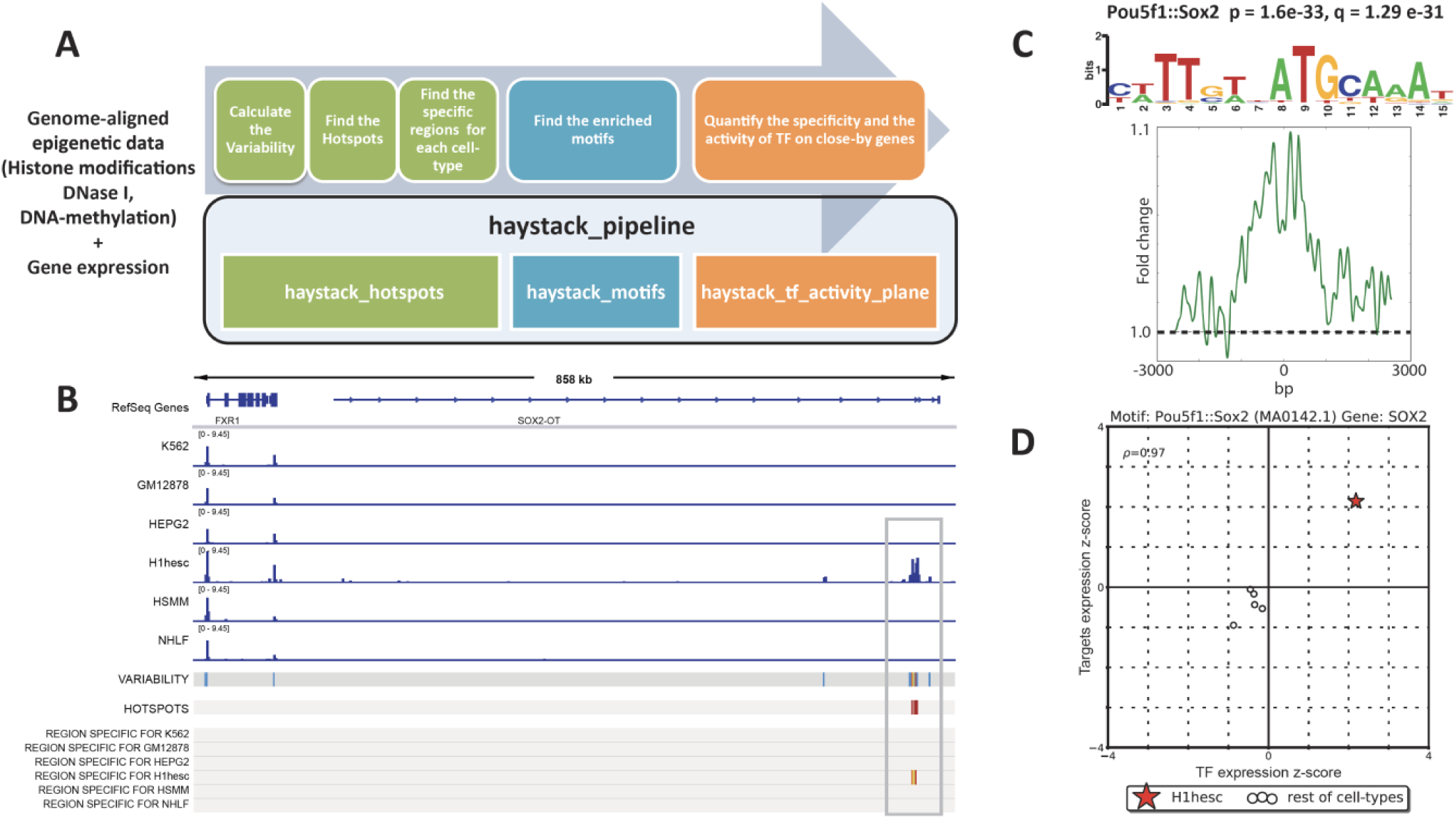
(A) Haystack overview: modules and corresponding functions. (B) Hotspot analysis on H3k27ac: signal tracks, variability track and hotspots of variability are computed from the ChIP-seq aligned data; the regions specific for a given cell type are also extracted. (C) Motif analysis on the regions specific for the H1hesc cell line: Pou5f1::Sox2 is significant; p-value and q-value, motif logo and average profile are calculated. (D) Transcription factor activity for Sox2 in H1esc (star) compared to the other cell types (circles), x-axis specificity of Sox2 expression (z-score), y-axis effect (z-score) on the gene nearby the regions containing the Sox2 motif.

### Module 1. Discovery of hotspots and cell-type specific regions

*haystack_hotspots* identifies the hotspots of variability, i.e., those regions that are highly variable for a given epigenetic mark among different cell types. The algorithm for identifying the hotspots was described previously in (Pinello, *et al*., 2014). Briefly the input for the pipeline is a set of genome-aligned sequencing tracks for a given epigenetic mark in different cell types, in BAM or bigWig format. The haystack_hotspots module first quantifies the sequence reads to nonoverlapping bins of predetermined size (500bp by default), and normalizes the data both using a variance stabilization method and quantile normalization. It then quantifies the variability of the processed data signal in each bin using the variance-to-mean ratio. The most variable regions, accordingly to this measure, are selected as hotspots (originally termed as Highly Plastic Regions in (Pinello, *et al*., 2014). The subsets of hotspot regions that have specific activity in a particular cell type are next identified, based on a z-score metric. Finally, an IGV (http://www.broadinstitute.org/igv/) XML session file is created to enable easy visualization of the results (Fig. 1B, Fig. S1).

### Module 2. Analysis of transcription factor motif

*haystack_motifs* identifies transcription factors (TFs) whose binding sequence motifs are enriched in a cell-type specific subset of hotspots. This module takes the output of *haystack_hotspots* as its input. Alternatively, the input may be a generic set of genomics regions; for example, promoters for a set of genes of interest or cell type specific enhancers. A motif database can also be specified (JASPAR (Mathelier, *et al*., 2016) by default) to look for motif enrichment (the basic counting of each motif is based on the FIMO software (Grant, *et al*., 2011)), with use of random or C+G content matched genomic sequences as background. We find that the latter option is more appropriate for histone modifications. The output of this module consists of an HTML page (Fig. S2) that reports each enriched motif, a series of informative parameters including the target/background ratio, the p-value (calculated with the Fisher’s exact test) and q-value, the motif logo, the central enrichment score, the average profile in the target regions containing the motif, and the closest genes for each region (Fig. 1C, Fig. S2).

### Module 3. Integration of gene expression data

Because different TFs may share similar sequence binding patterns, the exact regulator cannot be determined by motif enrichment analysis alone. Spurious association may also occur due to, for example, over-abundance of motif sequences. *haystack_tf_activity_plane* provides an additional filter to select for the most relevant TFs by further integrating gene expression data; it is based on the assumption that the expression level of a functional transcription factor is correlated with the expression level of the target genes of hotspot regions. Such a relationship is visualized with the use of an activity plane representation (Fig. 1D). Only those TF gene/motif pairs that show significant separation are reported in the final output (Fig. S3). Earlier we showed that such a filter is important for identifying factors that truly play a key role in mediating poised enhancer activities (Pinello, *et al*., 2014).

## 3 Results

### Analysis of H3K27ac data

To demonstrate Haystack’s utility, we analyzed 6 ChIP-seq datasets from the ENCODE project (Consortium, 2012) for the histone modification H3K27ac (Fig 1B). H3K27ac often marks active enhancers that promote the expression of nearby genes. We also integrated six RNA-seq assays, to quantify gene expression for the same cell types. Figure 1 shows the output of the pipeline: Haystack not only recovers regions that are highly dynamic (variability and hotspots tracks in Fig. 1), but also regions that are specifically active in each cell type. Additionally, Haystack detects several TFs that are likely to play an important regulatory role in those regions (Fig. S3). For example, for regions that are specifically active in the embryonic stem cell line (H1hesc), we found that the Pou5f1::Sox2 composed motif was highly enriched, and the expression of Sox2 a fundamental TF for embryonic stem cell identity was highly specific and positively correlated with activity of the target genes.

### Analysis of Roadmap Epigenomics Project

We applied the Haystack pipeline to data from the Roadmap Epigenomics Project using the maximal number of non-redundant cell-types for which gene expression and epigenetic data was available (Supplementary Section 4). We provide precomputed analysis for H3k27ac (41 cell types), H3K27me3 (41 cell types), H3K4me3 (41 cell types), and DNase I hypersensitivity (25 cell types). This analysis provides a valuable resource for researchers interested in in identifying functional elements in the human genome, exploring how epigenetic variability is controlled in different cell types and uncovering which sequence features play a role in gene regulation.

### Reproducible results through Cloud and Docker support

To facilitate the use of Haystack by prospective users without access to a computational facility, we provide detailed instructions in the supplementary materials on how to deploy and test Haystack on the Amazon Web Services cloud or similar services. We also provide a Docker image to enable users to easily use our tool and reproduce the analysis presented in this manuscript on any platform (see Supplementary Section 5).

## Acknowledgements

We would like to thank Stuart Orkin, Jian Xu, Nadin Rohland, Kimberly Glass, Eugenio Marco Rubio, Jialiang Huang, Assieh Saadatpour, Jennifer Wu, and Kendell Clement for their helpful discussions and/or for testing the software. This work was supported by National Institutes of Health award R00HG008399 (to L.P.) and R01HG009663 (to G-C Y.).

## Supplementary Information

### 1 Installation

The Haystack pipeline is implemented as a Python package called *haystack_bio*. The software has been tested on several operating systems: CentOS 6.5, Ubuntu 14.04 LTS, Ubuntu 16.04 LTS, OS X >=10.11. Although *haystack_bio* supports only 64-bit Linux and macOS, it can be run on Windows systems or other operating systems using a provided Docker image. For instructions on how to install the Docker software and the docker image, please refer to Section 5.1.

#### 1.1 Bioconda installation for Linux and macOS

*haystack_bio* and all its dependencies can be easily installed automatically through Bioconda (<u>https://bioconda.github.io/</u>), a software repository channel for the *conda* package manager. Bioconda streamlines the process of building and installing any software dependency that a package requires.

The entire installation process consists of three steps: installing Miniconda, adding the Bioconda channel, and installing *haystack_bio.* For the following steps, we are assuming you are on a 64-bit Linux or a macOS system and that you have not installed Miniconda/Anaconda before on your system. If you have *conda* already installed please skip to Step 3. To use *haystack_bio* on a Windows system please refer to the Docker section.

*Step 1*: Download the latest Miniconda 2 (Python 2.7) to your home directory.

For Linux:

~~~
      *wget -c https://repo.continuum.io/miniconda/Miniconda2-latest-Linux-x86_64.sh -O
      $HOME/miniconda.sh*
~~~

For macOS:

~~~
      *wget -c https://repo.continuum.io/miniconda/Miniconda2-latest-MacOSX-x86_64.sh -O
      $HOME/miniconda.sh*
~~~

*Step 2*: Execute the Miniconda setup file.

~~~
      *bash $HOME/miniconda.sh*
~~~

*Step 3*: Update the *conda* repository to include Bioconda by running these four lines in the order shown.

~~~
      *conda update --all --yes
      conda config --add channels defaults
      conda config --add channels conda-forge
      conda config --add channels bioconda*
~~~

*Step 4*: Install the *haystack_bio* package and its dependencies by simply running

~~~
      *conda install haystack_bio*
~~~

**Notes on installation:** For other installation options, please consult the Section 5. If you encounter any difficulty in installing Miniconda or adding the Bioconda channel, please refer to the Bioconda project’s website (<u>https://bioconda.github.io/</u>) for more detailed installation instructions. Note that if you have Miniconda/Anaconda 3 already installed, you would need to create a separate environment for Python 2.7. Please consult the conda.io webpage (<u>https://conda.io/docs/py2or3.html#create-python-2-or-3-environments</u>) for details.

#### 1.2 Testing the installation

To test if the package was successfully installed, please run the following command:

~~~
      *haystack_hotspots -h*
~~~

The *-h* help flag outputs a list of all the possible command line options that you can supply to the *haystack_hotspots* module. If you see such a list, then the software has been installed successfully.

We strongly suggest testing the entire pipeline on your system using the sample data for the hg19 reference genome bundled with the package by running the command

~~~
      *haystack_run_test*
~~~

The command will automatically download the hg19 reference genome. Since it is necessary to download about 800MB, this step could take a long time on a slow internet connection. We do not include any reference genome with the installation since those files tend to be rather big. If the test completes successfully, you should see the message “*Test completed successfully*” in the console.

The reference genome hg19 (or any other genome such as mm9, mm10, hg38) can be also downloaded before performing the test with the command

~~~
      *haystack_download_genome hg19*
~~~

The reference genomes and some required additional files are always saved into a dedicated genomes folder, by default in this path:

~~~
      *$HOME/miniconda/lib/python2.7/site-packages/haystack/haystack_data/genomes*
~~~

Note: The path could be slightly different depending on the Anaconda/Miniconda version installed (e.g. *$HOME/`miniconda2`*). Also note that since the sample test data included in the package is for 4 cell types only and is limited to a small fraction of the genome (i.e. chromosome 21), the motif analysis step in the pipeline might not return any positive findings for some cell types. The test run output can be found in the folder*$HOME/haystack_test_output*.

### 2 How to use haystack_bio

*haystack_bio* consists of the following three modules:

1. *haystack_hotspots*: finds regions that are variable across different ChIP-seq, DNase-seq, ATAC-seq or similar assays.
2. *haystack_motifs*: finds enriched transcription factor motifs in a given set of genomic regions.
3. *haystack_tf_activity_plane*: quantifies the specificity and the activity of the TFs highlighted by the *haystack_motif* integrating gene expression data

The command *haystack_pipeline* executes the whole pipeline automatically. That is, it executes Module 1 followed by Module 2 and Module 3 (optionally, if gene expression files are provided), finding hotspots, specific regions, motifs and quantifying their activity on nearby genes.

### 3 A walk-through example using ENCODE ChIP-seq data for the H3K27ac histone mark in six cell types

In this section, we showcase the commands using the example we have in the paper. We recreate the output produced by the pipeline when running on the entire genome with six cell types.

*Step 1*: Open a command terminal in the directory of your choosing and download the complete set of data files as an archive file to that directory.

~~~
      *wget-O data_h3k27ac_6cells.zip
      https://www.dropbox.com/s/4yjx7ypj0c82ryh/data_h3k27ac_6cells.zip?dl=1*
~~~

*Step 2*: Decompress the archive file

~~~
      *unzip data_h3k27ac_6cells.zip -d $HOME*
~~~

The command will unzip the data files archive into a folder called *`data_h3k27ac_6cells.`* Inside the folder you will see a *samples_names.txt* file containing the relative paths to the data files.

The file is a tab delimited text file with three columns containing

1. The sample name (cell type).
2. The path of the corresponding bam file.
3. The path of the gene expression file. Note that this last column is optional.

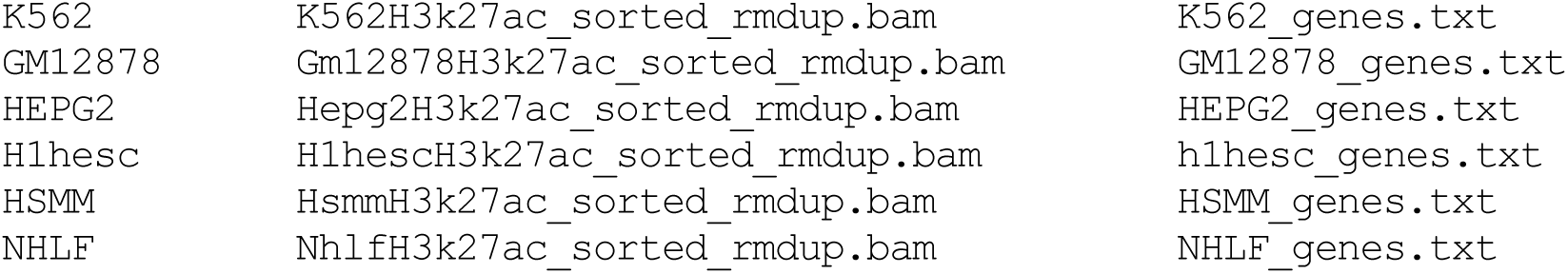

*Step 3*: Change directory and run the pipeline:

~~~
      *cd $HOME/data_h3k27ac_6cells
      haystack_pipeline samples_names.txt hg19 --output_directory
      $HOME/HAYSTACK_H3K27ac --blacklist hg19*
~~~

The *haystack_pipeline* command saves the output to the folder *`HAYSTACK_H3K27ac`* in your home directory. All the results will be stored in inside the sub-folder HAYSTACK_PIPELINE_RESULT. This will recreate the panels and the plots showed in Figure 1, plus other panels and plots for all the other cell types contained in the test dataset. The *`--blacklist`* flag accepts a file of blacklisted genomic regions in BED format. We suggest excluding those regions since they are characterized by artifact signals. For hg19, we have provided a BED file of blacklisted regions inside the package and this can be automatically loaded specifying just the string *`hg19`* as in our example. This file was obtained merging the ENCODE Data Analysis Consortium (DAC) Blacklisted Regions, the Duke Excluded Regions, and gap locations such as centromeres, telomeres, and contigs into one file.

The *haystack_pipeline* command is equivalent to running *haystack_hotspots* followed by *haystack_motifs* and *haystack_tf_activity_plane*.

You can run the pipeline also without creating a *samples_names.txt* file by providing the folder containing the *BAM* or *bigWig* files with the commands:

- Folder with BAM files: *`haystack_hotspots $HOME/data_h3k27ac_6cells hg19 --output_directory`* *`$HOME/HAYSTACK_H3K27ac --blacklist hg19`*
- Folder with bigwig files: *`haystack_hotspots $HOME/data_h3k27ac_6cells hg19 --output_directory`* *`$HOME/HAYSTACK_H3K27ac --blacklist hg19 --input_is_bigwig`*

Note, however, that in this case the pipeline runs *haystack_hotspots* and *haystack_motifs*, but not *haystack_tf_activity_plane* since no gene expression data are provided.

The inputs and outputs of the three modules of the pipeline are as follows.

### Module 1: haystack_hotspots

This step finds hotspots, which we define as regions that are highly variable across different cell types for a given epigenetic mark.

Sub-command run by *haystack_pipeline*:

~~~
      *haystack_hotspots $HOME/data_h3k27ac_6cells samples_names.txt hg19 --
      output_directory $HOME/HAYSTACK_H3K27ac --blacklist hg19*
~~~

Input:

- A tab delimited file with two columns: (1) the sample name and (2) the full path of the corresponding BAM or b*igWig* file.

~~~
K562         K562H3k27ac_sorted_rmdup.bam
GM12878      Gm12878H3k27ac_sorted_rmdup.bam
HEPG2        Hepg2H3k27ac_sorted_rmdup.bam
H1hesc       H1hescH3k27ac_sorted_rmdup.bam
HSMM         HsmmH3k27ac_sorted_rmdup.bam
NHLF         NhlfH3k27ac_sorted_rmdup.bam
~~~
- The reference genome (e.g. hg19)
- An optional bed file with blacklisted regions (the one for hg19 has been provided)

Output:

- The normalized *bigWig* files for each of the six samples. By default, we use quantile normalization. Please see the list of parameters in Section 9 for more details on the available normalization steps.
- A file of specific regions for each of the six samples. These are regions in which the signal is more enriched for a particular cell type compared to the rest.
- A file of background regions for each of the six samples.
- A *SELECTED_VARIABILITY_HOTSPOT*.bedgraph file containing the hotspots (i.e. regions that are most variable)
- An XML session file for the IGV software (http://www.broadinstitute.org/igv/) from the Broad Institute to easily visualize all the tracks produced, the hotspots and the specific regions for each cell line. To load it, just drag and drop the file OPEN_ME_WITH_IGV.xml_ from the output folder on top of the IGV window or alternatively load it in IGV with File-> Open Session…If you have trouble opening the file please update your IGV version. Additionally, please do not move the XML file only since you need all the files in the output folder to correctly load the session.

**Figure S1.**
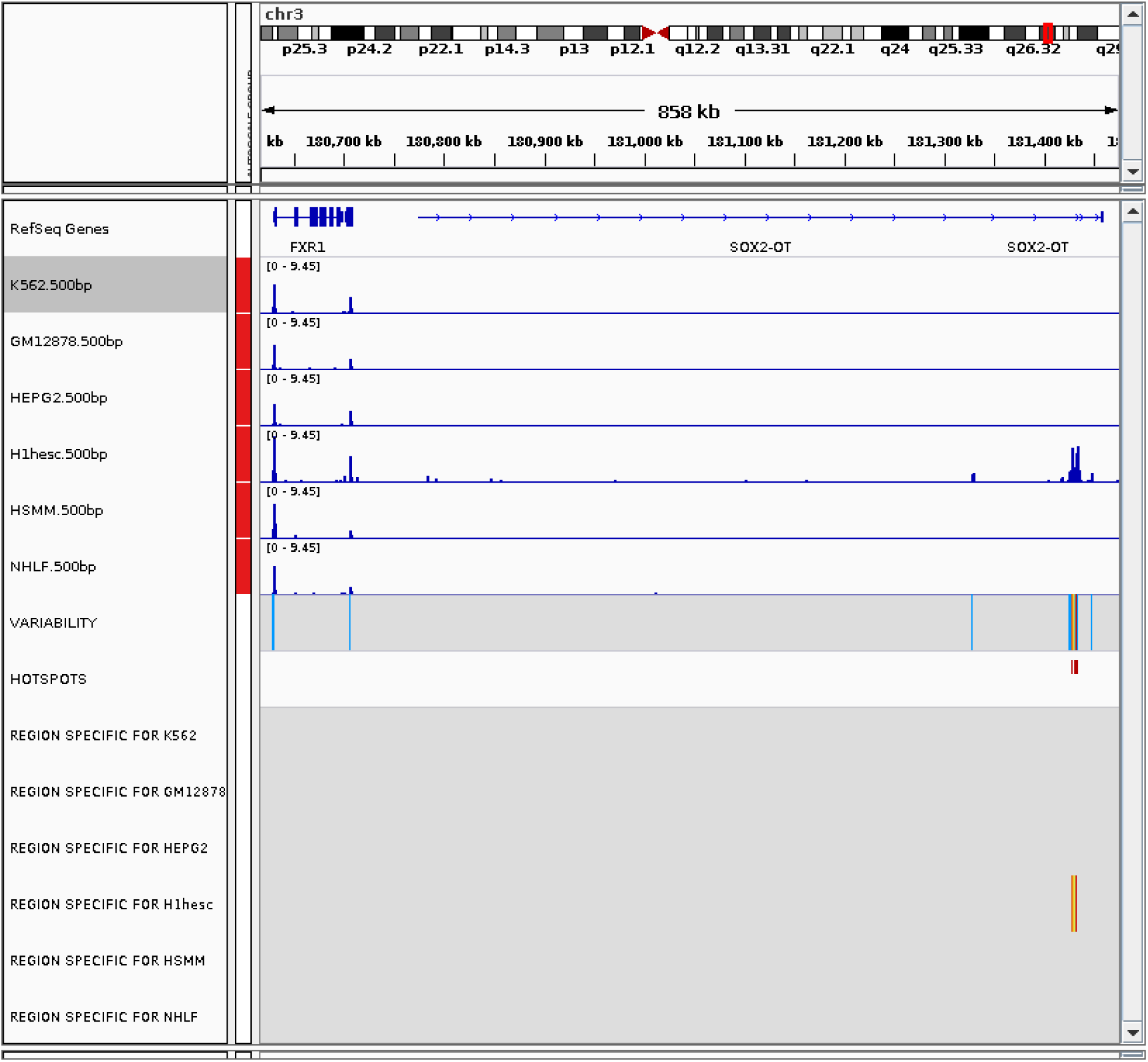
Screenshot of the IGV browser showing the bigwig tracks, the hotspots, and the specific regions.

Notes:

- IMPORTANT: Folder names and file paths should not have white spaces. Please use underscore instead.
- If you are running haystack_hotspots using bigWig files you need to add the option: --input_is_bigwig
- The haystack_download_genome command allows you to download and add a reference genome from UCSC to haystack_bio in the appropriate format. To download a particular genome run:

~~~
haystack_download_genome genome_name
~~~ You might not need to call this command explicitly since it is called automatically when you run the pipeline.

### Module 2: haystack_motifs

This step finds enriched transcription factor motifs in a given set of genomic regions. When executed from the pipeline it automatically analyzes cell type specific genomic regions obtained from Module 1 and identifies transcription factors whose binding sequence motifs are enriched in those regions.

Sub-command run by *haystack_pipeline* (for each sample):

~~~
      *haystack_motifs specific_regions_filename hg19 --bed_bg_filename
      background_regions_filename --name sample_name --output_directory
      $HOME/HAYSTACK_H3K27ac/HAYSTACK_PIPELINE_RESULT/HAYSTACK_MOTIFS*
~~~

Input:

- Genomic regions file (i.e. *Regions_specific_for_K562.500bp_z_1.50.bed*, or a list of enhancer or promoter regions)
- The reference genome (i.e. hg19)
- Background regions file (i.e. *Background_for_K562.500bp_z_0.25.bed*)
- Sample name (i.e. K562)

Output:

The output consists of an HTML report for each sample.

Each row in the table corresponds to an enriched motif. There are 12 columns in the table corresponding to following variables.

- Motif id
- Motif name
- Presence in Target: Frequency of its presence in the specific region file of the corresponding sample
- Presence in BG: Frequency of its presence in the background region file of the corresponding sample
- The ratio of Presence in Target to Presence in BG
- The p value (calculated with the Fisher’s exact test)
- The q values: corrected p-values using BH
- Central enrichment: ratio between the average signal in central part of the profile and the flanking sides (by default the central region is defined as 1/5 of the entire window)
- Motif profile: average presence of the motifs inside the regions profiles
- Motif logo: Logo obtained with Weblogo
- List of regions with motifs and coordinates of the motifs in those regions
- List of closest genes to the regions with a particular motif

**Figure S2.**
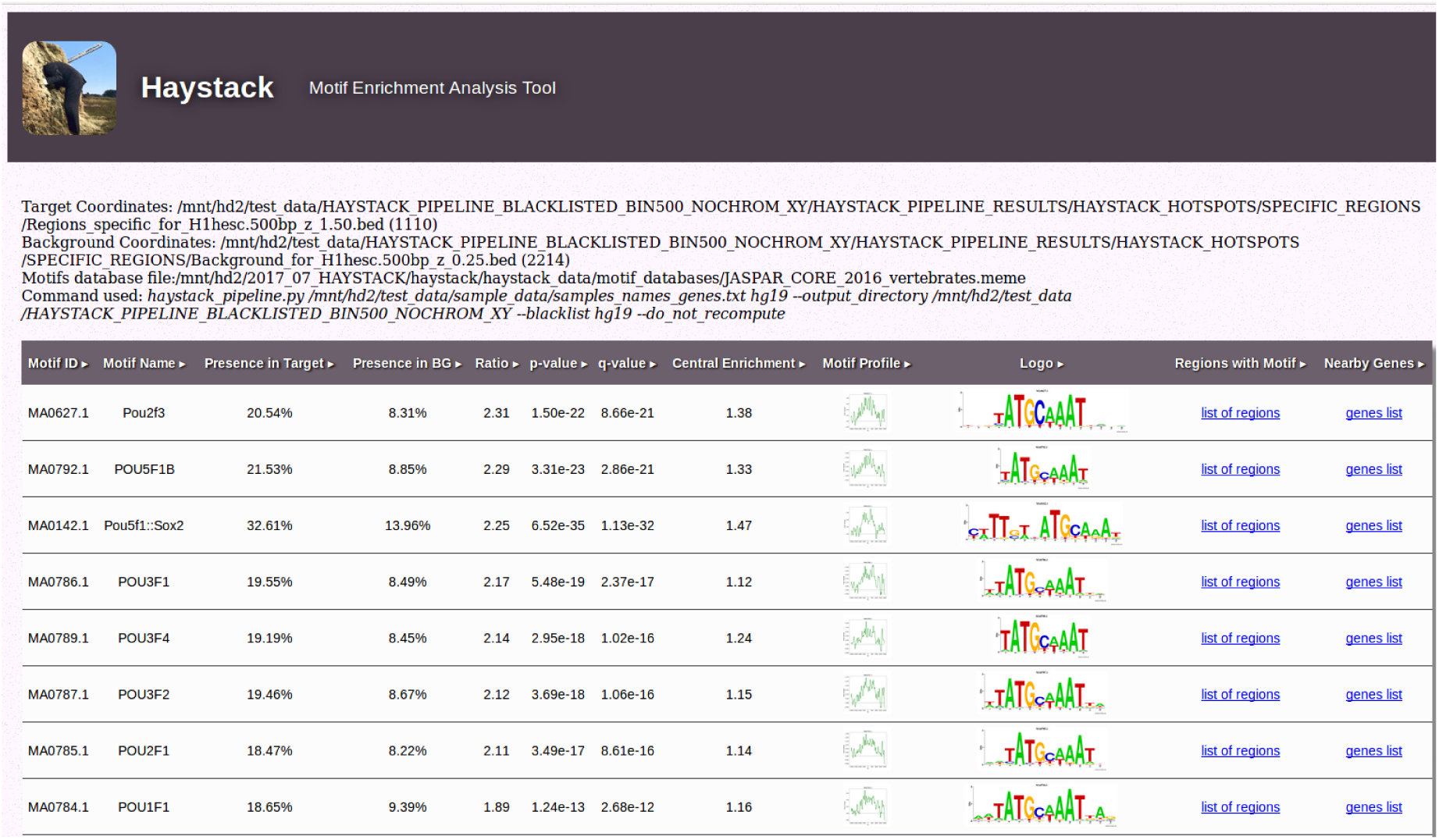
Screenshot of the HTML report generated for the H1hesc sample.

**Notes:**

- It is possible to run the motif analysis only by calling the *haystack_motifs* module on a given set of genomic regions. For example, to analyze the *BED* file *myregions.bed* on the hg19 genome, run

~~~
*haystack_motifs myregions.bed hg19*
~~~
- To specify a custom background file for the analysis, for example *mybackgroundregions.bed* run

~~~
*haystack_motifs myregions.bed hg19 --bed_bg_filename mybackgroundregions.bed*
~~~
- To use a particular motif database (the default is JASPAR) use

~~~
*haystack_motifs myregions.bed hg19 --meme_motifs_filename my_database.meme*
~~~
- Note that the database file must be in the MEME format: <u>http://meme.nbcr.net/meme/doc/meme</u>format.html#min_format

### Module 3: haystack_tf_activity_plane

This step acts as an additional filter to restrict the set of enriched transcription factors found by Module 2 to those that are specifically expressed in the cell of interest and that show a positive or negative correlation with expression of genes nearby their binding sites (see Figure S3).

Sub-command run by *haystack_pipeline* (for each sample):

~~~
      *haystack_tf_activity_plane
      $HOME/HAYSTACK_H3K27ac/HAYSTACK_PIPELINE_RESULT/HAYSTACK_MOTIFS
      sample_names_tf_activity_filename sample_name --output_directory
      $HOME/HAYSTACK_H3K27ac/HAYSTACK_PIPELINE_RESULT/HAYSTACK_TFs_ACTIVITY_PLANES*
~~~

Input:

- An output folder generated by *haystack_motif*
- A tab delimited file describing the samples names and the gene expression filenames to use (for example the first and third column of *`samples_names.txt`* used by the pipeline):

~~~
        K562      K562_genes.txt
        GM12878   GM12878_genes.txt
        HEPG2     HEPG2_genes.txt
        H1hesc    h1hesc_genes.txt
        HSMM      HSMM_genes.txt
        NHLF      NHLF_genes.txt
~~~
- A set of files containing gene expression data specified in a tab delimited format with two columns: (1) gene symbol and (2) gene expression value, for example the file *`K562_genes.txt`* may contain the lines:

~~~
        RNF14 7.408579
        UBE2Q1 9.107306
        UBE2Q2 7.847002
        RNF10 9.500193
        RNF11 7.545264
        LRRC31 3.477048
        RNF13 7.670409
        CBX4 7.070998
        REM1 6.148991
        .
        .
        .
~~~

Output:

- A set of figures each containing the TF activity plane for a given motif.

**Figure S3.**
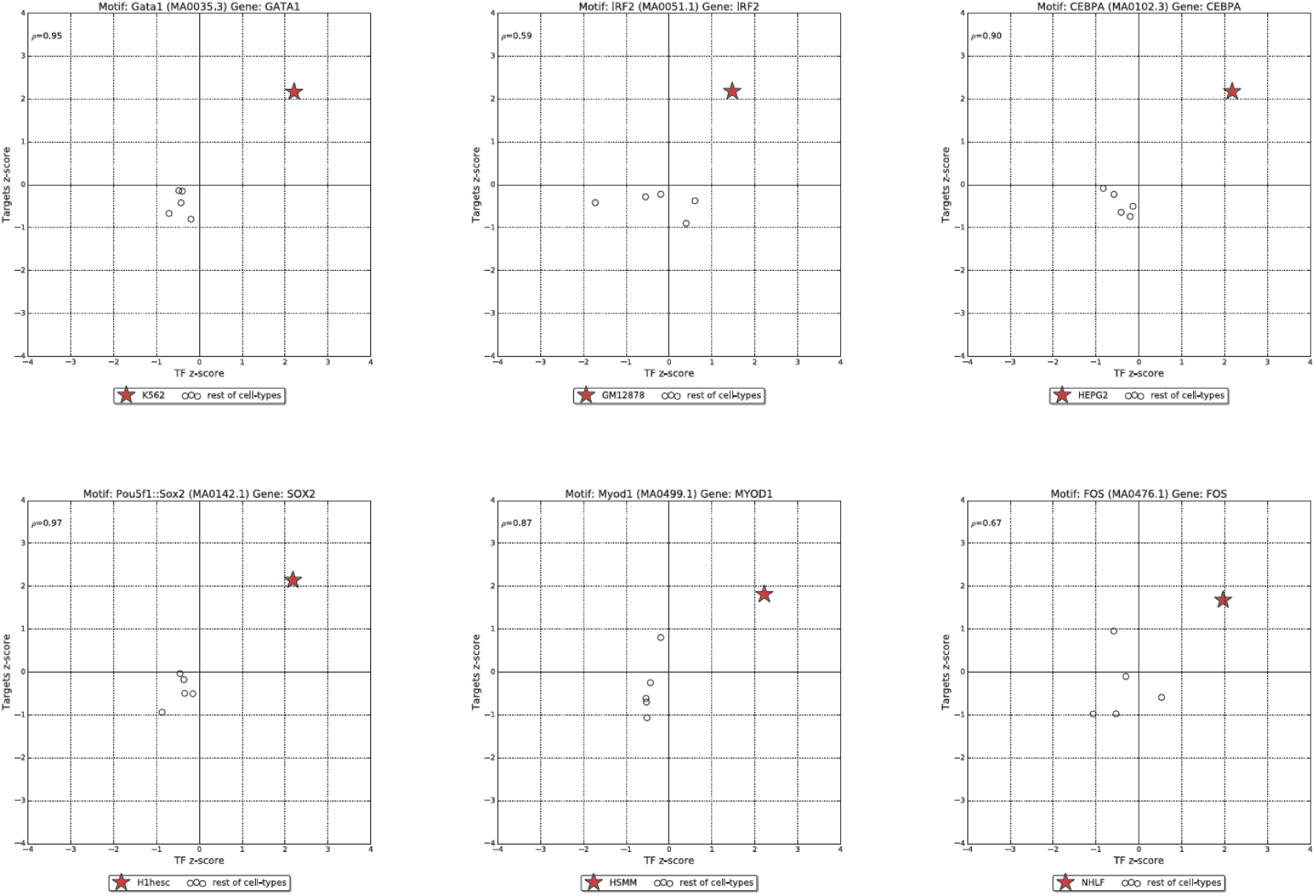
Activity planes corresponding to enriched motif for each of the six samples (cell types). Each sub-figure depicts the relationship between the expression of a transcription factor and the expression level of the target genes of cell type specific regions hotspot regions.

### 4 Analysis of Roadmap Epigenomic Project Data

We reanalyzed data from the Roadmap Epigenomics Project using the Haystack pipeline. We extended the example in Section 3.2 (limited to 6 cell types only for illustration purposes) to 41 cell lines, the maximal number of nonredundant cell-types for which gene expression and H3K27ac data are available. We also made available precomputed results for three other epigenetic marks: H3K27me3 (41 cell types), H3K4me3 (41 cell types), and DNase I hypersensitivity (25 cell types). Please see Supplementary Materials Section 4 for additional details regarding the systematic analysis of Roadmap Epigenomics data for these 4 epigenomic marks. Given that the Haystack pipeline provides IGV tracks and a session file with the summary of the analysis, the interested user just needs to open the session file (an XML file) included in the output folder with the IGV software to explore the precomputed analysis (Figure S4), and then investigate any region or gene of interest (Figure S5). Researchers will not only see the variable regions, cell-type specific regions, and input signals together, but will also see a gene annotation track (the default is RefSeq) or any of the numerous other annotations available for the IGV software.

Moreover, because the output tracks conform to an open and standard format (bigWig file for signal tracks and BED file for genomic regions), it is also possible to load the tracks and regions produced by Haystack into the UCSC Genome Browser, or other software, for further data integration. For the motif and activity plane analysis (Figure S6), we provide HTML reports and PDF files with logos, p/q-value and profiles for each cell type included in the analysis, with links to the relevant files for more detailed analysis.

Precomputed results for four epigenetic marks can be downloaded using the following links.

- H3K27ac (41 cell types): <u>https://www.dropbox.com/s/30vcy2kasv5azp6/haystack_H3K27ac_41cells.zip?dl=1</u>
- H3K27me3 (41 cell types): <u>https://www.dropbox.com/s/pwvy7lum0d4iums/haystack_H3K27me3_41cells.zip?dl=1</u>
- H3K4me3 (41 cell types): <u>https://www.dropbox.com/s/mgvwb9mepbqn6tz/haystack_H3K4me3_41cells.zip?dl=1</u>
- DNase I (25 cell types): <u>https://www.dropbox.com/s/51wsk91zg01sx3y/haystack_DNase_25cells.zip?dl=1</u>

**Figure S4.**
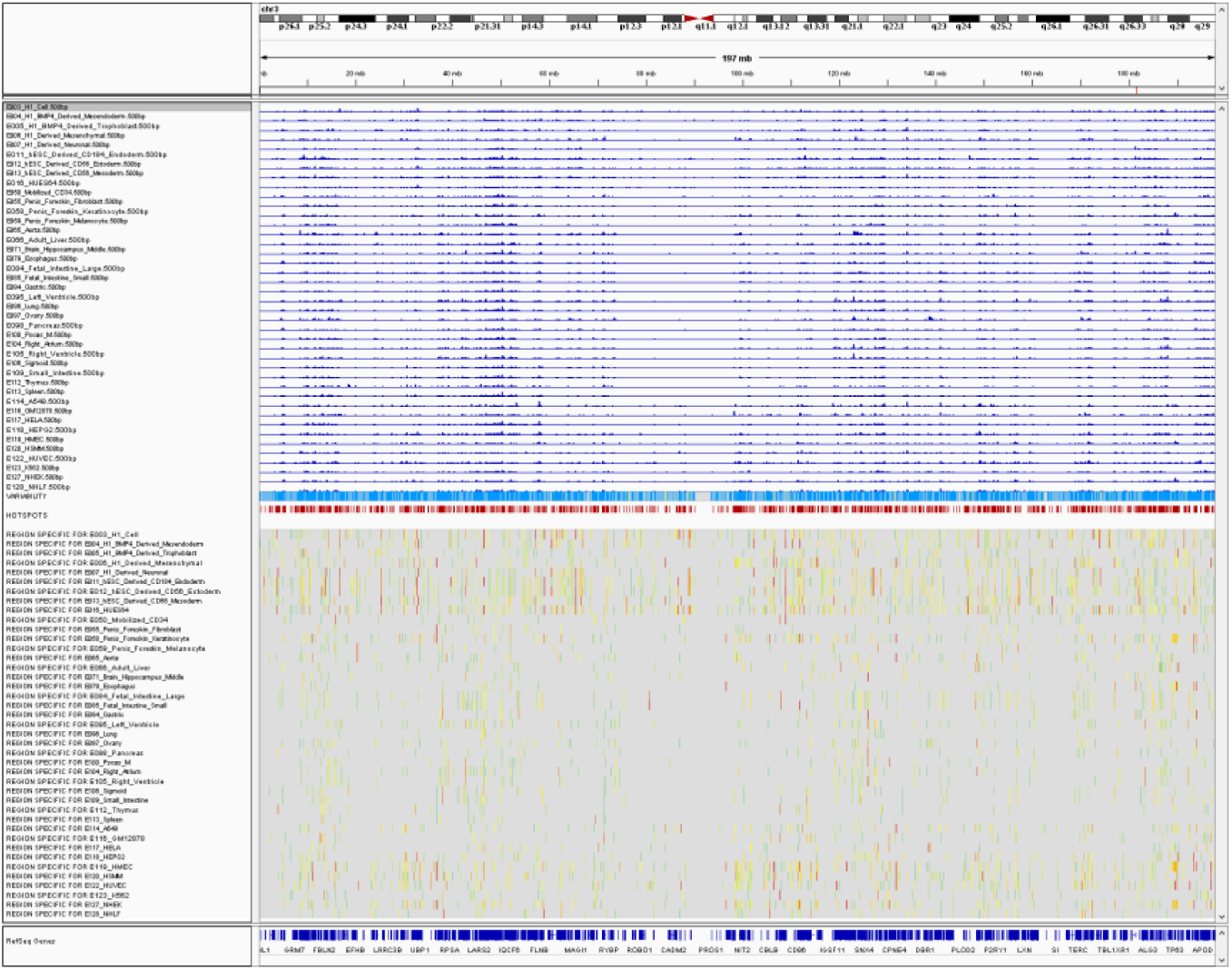
Screenshot of IGV browser showing the bigwig tracks, the hotspots, and the specific regions for Chromosome 3.

**Figure S5.**
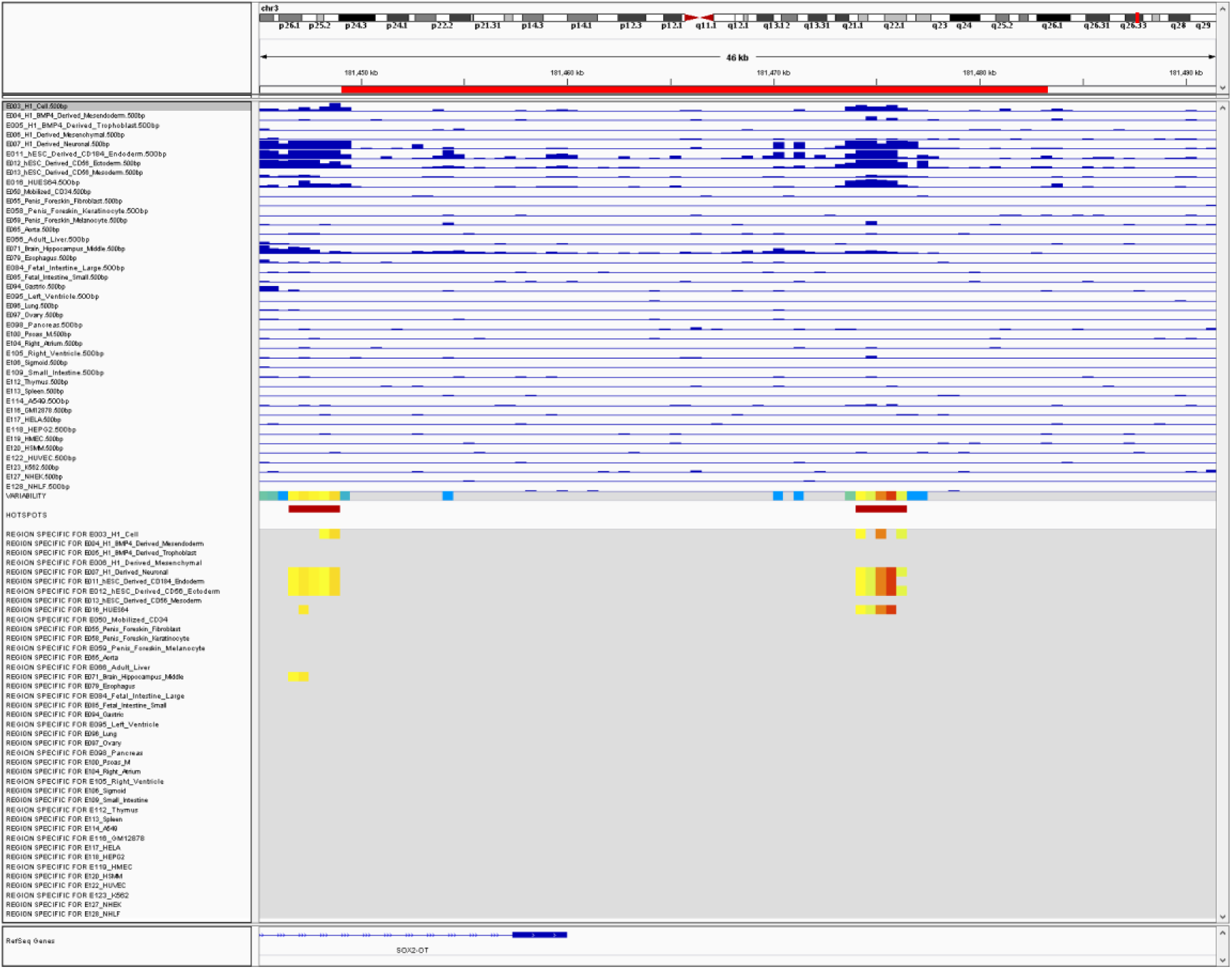
Screenshot of IGV browser showing the bigwig tracks, the hotspots, and the specific regions for Sox2.

**Figure S6.**
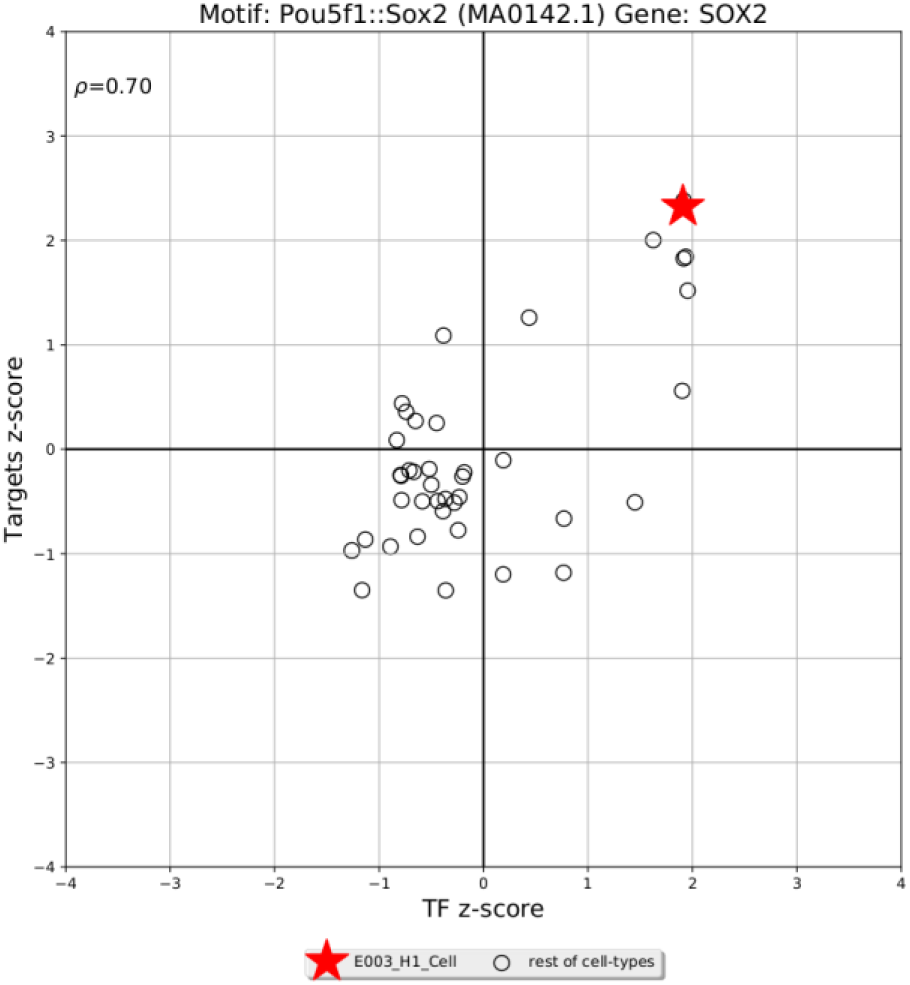
Transcription factor activity for Sox2 in H1esc (star) compared to the other cell types (circles), x-axis specificity of Sox2 expression (z-score), y-axis effect (z-score) on the gene nearby the regions containing the Sox2 motif.

We also provide R scripts and notebook showing how to accomplish several tasks you would probably need if you will be working with large amount of data from the ENCODE, Roadmap Epigenomics projects, or other consortia. These scripts can help you in the following tasks:

- Downloading epigenomic data for any experiment and cell type.
- Determining which combination of histone mark (experiment) are present across a given set of cell types.
- Preprocessing gene expression data and converting gene ids from one form to another (Ensembl ID).
- Creating *sample_names* text files with the data file paths for the haystack pipeline.

The scripts and the interactive notebook can be found in the project’s GitHub repository folder

~~~
*
        haystack_bio/scripts/roadmap_data_scripts*
~~~

### 5 Other Installation Options

#### 5.1 Docker image

*haystack_bio* can be easily used without installation using our provided Docker image. Docker is a virtualization technology that allows the creation of reproducible and isolated environments for any software. You can install Docker by following the instruction for your platform at <u>https://www.docker.com/.</u>

Note: For Linux platforms, make sure to run the Docker post-installation instructions so you can run the command without *sudo* privileges.

After the installation is complete you can download the Docker image for *haystack_bio* by simply running

~~~
      *docker pull pinellolab/haystack_bio*
~~~

Before using the *haystack_bio image*, first create a *haystack_genomes* folder in your home directory to keep a persistent copy of the genomes you will be downloading:

*`mkdir ${HOME}/haystack_genomes`*

This allows the use of the *-v* option to link the <u>full path</u> of the *haystack_genomes* folder you have created on your host to the *haystack_genomes* folder used inside the *haystack_bio* container. You need also to mount the data folder containing the files you are going to use with an additional -v option. For example, assuming you have the *samples_names.txt* and the BAM files listed in it the current folder and the current folder is called *`data_h3k27ac_6cells`*, you can use the following command:

~~~
      *docker run -v ${PWD}:/docker_data \
               -v ${HOME}/haystack_genomes:/haystack_genomes \
               -w /docker_data -it pinellolab/haystack_bio haystack_pipeline \
               samples_names.txt hg19 --blacklist hg19*
~~~

If you run Docker on Window you should specify the full path of the data as such:

~~~
      *docker run -v //c/Users/Username/Downloads/data_h3k27ac_6cells:/docker_data -
      v //c/Users/Username/haystack_genomes:/haystack_genomes -w /docker_data -it
      pinelloalab/haystack_bio haystack_pipeline samples_names.txt hg19 --blacklist
      hg19*
~~~

Where *Username* is your account user name. Running other commands can be done with the same syntax and path conventions.

#### Allocation of memory for Docker containers

It might be necessary to manually increase the allocated memory to the container. The default memory assigned by Docker may depend on the version and on the machine upon which it is run. To run the *`haystack_bio`* container with the provided example, we suggest assigning at least 8GB of RAM to the docker container. For analysis involving many tracks, it may be necessary to increase the amount of allocated memory under the Settings…/ Advanced panel (see Figure S7).

**Figure S7.**
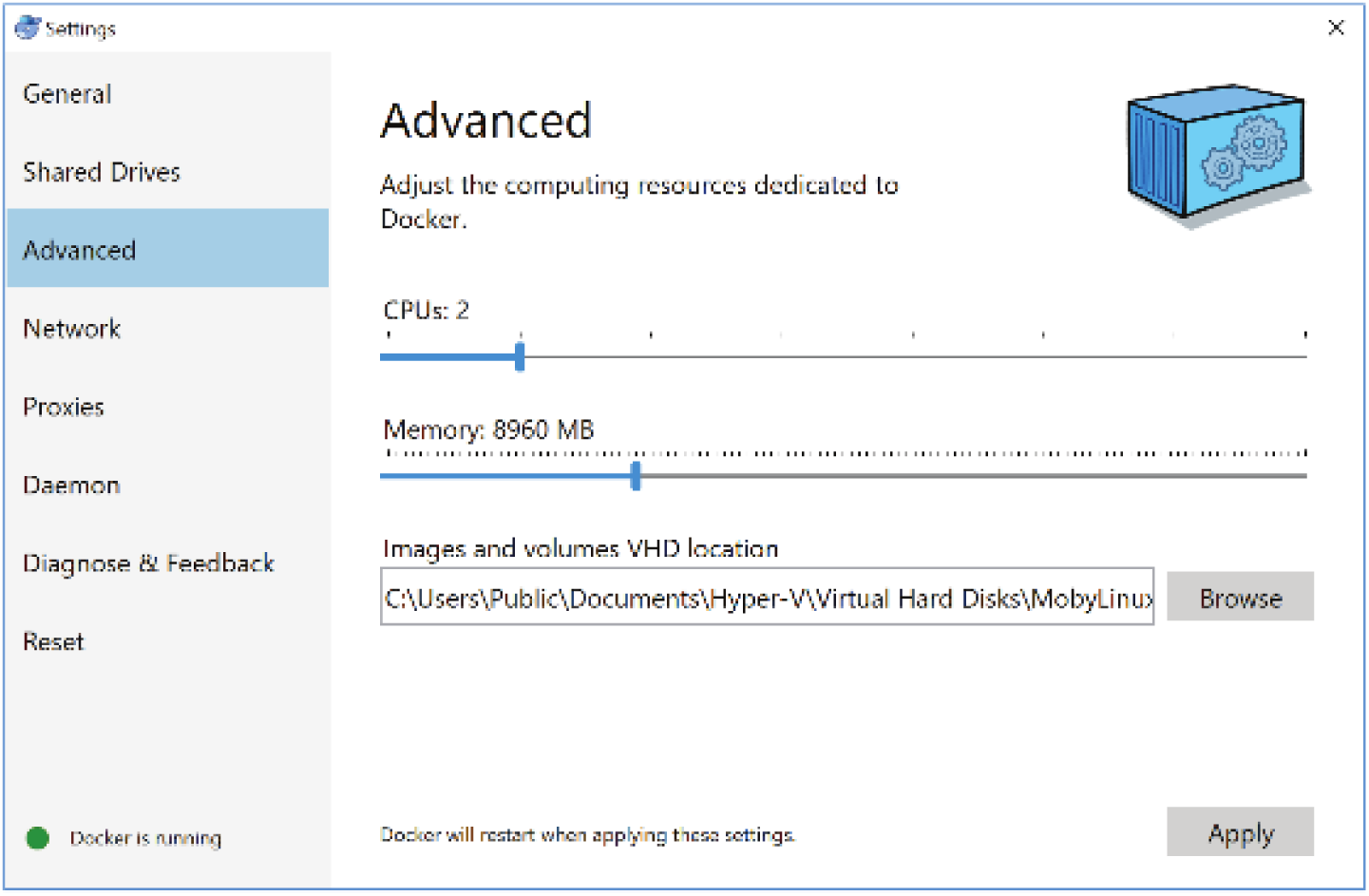
Screenshot of Docker’s settings tab.

#### 5.2 Advanced Installation on other platforms

First make sure that the following software and package dependencies are installed.

Software:

~~~
        *- ghostscript
        - meme 4.11.2
        - bedtools
        - sambamba
        - ucsc bigwigaverageoverbed
        - ucsc bedgraphtobigwig*
~~~

Python 2.7 packages:

~~~
        *- setuptools
        - bx-python
        - numpy
        - scipy
        - matplotlib
        - jinja2
        - pandas
        - tqdm
        - weblogo*
~~~

After installing all the dependencies, please download the repository and execute the command inside the root folder.

~~~
      *python setup.py install*
~~~

The Docker image recipe found in Dockerfile installs and builds the above-listed dependencies on Ubuntu 16.04. You can take this file as example and modify the build steps to manually install *haystack_bio* on other platforms.

#### Installation script for Ubuntu 16.04 on a local machine or on a cloud instance of Amazon Web Service (AWS)

We provide an installation script that downloads, builds, and installs haystack_bio and all its dependencies. To download and execute the file, run the following three commands.

~~~
      *wget-c*
      https://raw.githubusercontent.com/pinellolab/haystack_bio/master/scripts/manu
      *al_build.sh -O $HOME/manual_build.sh

      chmod +x $HOME/manual_build.sh

      $HOME/manual_build.sh -b*
~~~

After the script finishes its execution, edit the file named *.bashrc* in your home direction and add the following line:

~~~
      *export PATH=$HOME/haystack_bio/binaries:$HOME/haystack_bio/binaries/meme/bin:$PATH*
~~~

To install on AWS, first launch and connect to the Amazon Instance you have chosen from the AWS console (is suggested to use an m3.large) or to your Ubuntu machine. Second, create a swap partition.

~~~
      *sudo dd if=/dev/zero of=/mnt/swapfile bs=1M count=20096

      sudo chown root:root /mnt/swapfile

      sudo chmod 600 /mnt/swapfile

      sudo mkswap /mnt/swapfile

      sudo swapon /mnt/swapfile

      sudo sh -c “echo ’/mnt/swapfile swap swap defaults 0 0’ >> /etc/fstab”

      sudo swapon -a*
~~~

Then, download and execute the *`manual_build.sh`* script file.

### 6 Jupyter Analysis Notebook

We have provided an interactive analysis Jupyter notebook for the first module of the pipeline. You can gain a more detailed insight of this part of the pipeline or extend it by examining the code and output. The notebook can be accessed at this link

*<u>http://nbviewer.jupyter.org/github/pinellolab/haystack_bio/blob/master/haystack/scripts/haystack_jypyter_not</u> ebook.ipynb*

### 7 Contact

We use GitHub issues (<u>https://github.com/pinellolab/haystack_bio/issues</u>) for tracking requests and bugs. Please submit a new issue if you any comment or you would like to report a software bug.

### 8 Third party software included and used in this distribution

PeakAnnotator: http://www.ebi.ac.uk/research/bertone/software.

### 9 List of all the parameters for each module

For usage instructions, please add the help flag *-h* to the module’s command to display what parameters can be provided to the pipeline or the individual modules. For example,

~~~
      *haystack_pipeline.py -h*
~~~

would output the usage and parameters’ description for the entire pipeline

~~~
      *haystack_pipeline.py [-h] [--name NAME]
                                        [--output_directory OUTPUT_DIRECTORY]
                                        [--bin_size BIN_SIZE] [--do_not_recompute]
                                        [--depleted] [--input_is_bigwig]
                                        [--disable_quantile_normalization]
                                        [--transformation {angle,log2,none}]
                                        [--z_score_high Z_SCORE_HIGH]
                                        [--z_score_low Z_SCORE_LOW] [--th_rpm TH_RPM]
                                        [--meme_motifs_filename MEME_MOTIFS_FILENAME]
                                        [--motif_mapping_filename MOTIF_MAPPING_FILENAME]
                                        [--plot_all] [--keep_intermediate_files]
                                        [--n_processes N_PROCESSES]
                                        [--blacklist BLACKLIST]
                                        [--chrom_exclude CHROM_EXCLUDE]
                                        [--read_ext READ_EXT]
                                        [--temp_directory TEMP_DIRECTORY] [--version]
                                        samples_filename_or_bam_folder genome_name

positional arguments:
 samples_filename_or_bam_folder
                                A tab delimeted file with in each row (1) a sample
                                name, (2) the path to the corresponding bam filename,
                                (3 optional) the path to the corresponding gene
                                expression filename.
 genome_name                    Genome assembly to use from UCSC (for example hg19,
                                mm9, etc.)

optional arguments:
 -h, --help                     show this help message and exit
 --name NAME                     Define a custom output filename for the report
 --output_directory OUTPUT_DIRECTORY
                                Output directory (default: current directory)
 --bin_size BIN_SIZE            bin size to use (default: 500bp)
 --do_not_recompute             Keep any file previously precalculated
 --depleted                     Look for cell type specific regions with depletion of
                                signal instead of enrichment
 --input_is_bigwig              Use the bigwig format instead of the bam format for
                                the input. Note: The files must have extension .bw
 --disable_quantile_normalization
                                Disable quantile normalization (default: False)
 --transformation {angle,log2,none}
                                Variance stabilizing transformation among: none,
 log2,
                                angle (default: angle)
 --z_score_high Z_SCORE_HIGH
                                z-score value to select the specific regions
 (default:
                                1.5)
 --z_score_low Z_SCORE_LOW
                                z-score value to select the not specific
                                regions (default: 0.25)
 --th_rpm TH_RPM                Percentile on the signal intensity to consider for
the
                                hotspots (default: 99)
 --meme_motifs_filename MEME_MOTIFS_FILENAME
                                Motifs database in MEME format (default JASPAR CORE
                                2016)
 --motif_mapping_filename MOTIF_MAPPING_FILENAME
                                Custom motif to gene mapping file (the default is for
                                JASPAR CORE 2016 database)
 --plot_all                     Disable the filter on the TF activity and correlation
                                (default z-score TF>0 and rho>0.3)
 --keep_intermediate_files
                                keep intermediate bedgraph files
 --n_processes N_PROCESSES
                                Specify the number of processes to use. The default
 is
                                #cores available.
 --blacklist BLACKLIST
                                Exclude blacklisted regions. Blacklisted regions are
                                not excluded by default. Use hg19 to blacklist
 regions
                                for the human genome 19, otherwise provide the
                                filepath for a bed file with blacklisted regions.
 --chrom_exclude CHROM_EXCLUDE
                                Exclude chromosomes. For example (_|chrM|chrX|chrY).
 --read_ext READ_EXT            Read extension in bps (default: 200)
 --temp_directory TEMP_DIRECTORY
                                Directory to store temporary files (default: /tmp)
 --version                      Print version and exit.*
~~~

### Module 1: haystack_hotspots

~~~
*haystack_hotspots [-h]  [--output_directory OUTPUT_DIRECTORY]
                                [--bin_size BIN_SIZE] [--chrom_exclude CHROM_EXCLUDE]
                                [--th_rpm TH_RPM] [--transformation {angle,log2,none}]
                                [--z_score_high Z_SCORE_HIGH]
                                [--z_score_low Z_SCORE_LOW] [--read_ext READ_EXT]
                                [--max_regions_percentage MAX_REGIONS_PERCENTAGE]
                                [--name NAME] [--blacklist BLACKLIST] [--depleted]
                                [--disable_quantile_normalization]
                                [--do_not_recompute] [--input_is_bigwig]
                                [--keep_intermediate_files]
                                [--n_processes N_PROCESSES] [--version]
                                samples_filename_or_bam_folder genome_name

 positional arguments:
  samples_filename_or_bam_folder
                                A tab delimited file with in each row (1) a sample
                                name, (2) the path to the corresponding bam or bigwig
                                filename. Alternatively it is possible to specify a
                                folder containing some .bam files to analyze.
  genome_name                   Genome assembly to use from UCSC (for example hg19,
                                mm9, etc.)
 optional arguments:
 -h, --help                     show this help message and exit
 --output_directory OUTPUT_DIRECTORY
                                Output directory (default: current directory)
 --bin_size BIN_SIZE             bin size to use(default: 500bp)
 --chrom_exclude CHROM_EXCLUDE
                                Exclude chromosomes. For example (_|chrM|chrX|chrY).
 --th_rpm TH_RPM                Percentile on the signal intensity to consider for the hotspots (default: 99)
 --transformation {angle,log2,none}
                                Variance stabilizing transformation among: none, log2,
                                angle (default: angle)
 --z_score_high Z_SCORE_HIGH
                               z-score value to select the specific regions (default: 1.5)
 --z_score_low Z_SCORE_LOW
                               z-score value to select the not specific regions (default: 0.25)
 --read_ext READ_EXT           Read extension in bps (default: 200)
 --max_regions_percentage MAX_REGIONS_PERCENTAGE
                               Upper bound on the % of the regions selected (default:
                               0.1, 0.0=0% 1.0=100%)
 --name NAME                   Define a custom output filename for the report
 --blacklist BLACKLIST
                               Exclude blacklisted regions. Blacklisted regions are
                               not excluded by default. Use hg19 to blacklist regions
                               for the human genome build 19, otherwise provide the
                               filepath for a bed file with blacklisted regions.
 --depleted                    Look for cell type specific regions with depletion of
                               signal instead of enrichment
 --disable_quantile_normalization
                               Disable quantile normalization (default: False)
 --do_not_recompute            Keep any file previously precalculated
 --input_is_bigwig             Use the bigwig format instead of the bam format for
                               the input. Note: The files must have extension .bw
 --keep_intermediate_files
                               keep intermediate bedgraph files
 --n_processes N_PROCESSES
                               Specify the number of processes to use. The default is
                               #cores available.
 --version                     Print version and exit.*
~~~

### Module 2: haystack_motifs

~~~
*
 haystack_motifs.py [-h]         [--bed_bg_filename BED_BG_FILENAME]
                                 [--meme_motifs_filename MEME_MOTIFS_FILENAME]
                                 [--nucleotide_bg_filename NUCLEOTIDE_BG_FILENAME]
                                 [--p_value P_VALUE] [--no_c_g_correction]
                                 [--c_g_bins C_G_BINS] [--mask_repetitive]
                                 [--n_target_coordinates N_TARGET_COORDINATES]
                                 [--use_entire_bg] [--bed_score_column BED_SCORE_COLUMN]
                                 [--bg_target_ratio BG_TARGET_RATIO] [--bootstrap]
                                 [--temp_directory TEMP_DIRECTORY]
                                 [--no_random_sampling_target] [--name NAME]
                                 [--internal_window_length INTERNAL_WINDOW_LENGTH]
                                 [--window_length WINDOW_LENGTH]
                                 [--min_central_enrichment MIN_CENTRAL_ENRICHMENT]
                                 [--disable_ratio] [--dump]
                                 [--output_directory OUTPUT_DIRECTORY]
                                 [--smooth_size SMOOTH_SIZE]
                                 [--gene_annotations_filename GENE_ANNOTATIONS_FILENAME]
                                 [--gene_ids_to_names_filename
 GENE_IDS_TO_NAMES_FILENAME]
                                 [--n_processes N_PROCESSES] [--version]
                                 bed_target_filename genome_name

 positional arguments:
 bed_target_filename             A bed file containing the target coordinates on the
                                 genome of reference
 genome_name                     Genome assembly to use from UCSC (for example hg19,
                                 mm9, etc.)
 optional arguments:
 -h, --help                      show this help message and exit
 --bed_bg_filename BED_BG_FILENAME
                                A bed file containing the backround coordinates on the
                                genome of reference (default random sampled regions
                                from the genome)
 --meme_motifs_filename MEME_MOTIFS_FILENAME
                                Motifs database in MEME format (default JASPAR CORE 2016)
 --nucleotide_bg_filename NUCLEOTIDE_BG_FILENAME
                                Nucleotide probability for the background in MEME
                                format (default precomupted on the Genome)
 --p_value P_VALUE              FIMO p-value for calling a motif hit significant
                                (deafult: 1e-4)
 --no_c_g_correction            Disable the matching of the C+G density of the
                                background
 --c_g_bins C_G_BINS            Number of bins for the C+G density correction
                                (default: 8)
 --mask_repetitive              Mask repetitive sequences
 --n_target_coordinates         N_TARGET_COORDINATES
                                Number of target coordinates to use (default: all)

 --use_entire_bg                Use the entire background file (use only when the cg correction is disabled)
 --bed_score_column BED_SCORE_COLUMN
                                Column in the bedfile that represents the score (default: 5)
 --bg_target_ratio BG_TARGET_RATIO
                                Background size/Target size ratio (default: 1.0)
 --bootstrap                    Enable the bootstrap if the target set or the
                                background set are too small, choices: True, False (default: False)
 --temp_directory TEMP_DIRECTORY
                               Directory to store temporary files (default: /tmp)
 --no_random_sampling_target
                               Select the best --n_target_coordinates using the score
                               column from the target file instead of randomly select them
 --name NAME                   Define a custom output filename for the report
 --internal_window_length INTERNAL_WINDOW_LENGTH
                               Window length in bp for the enrichment (default:
                               average lenght of the target sequences)
 --window_length WINDOW_LENGTH
                               Window length in bp for the profiler (default:
                               internal_window_length*5)
 --min_central_enrichment MIN_CENTRAL_ENRICHMENT
                               Minimum central enrichment to report a motif
                               default:>1.0)
 --disable_ratio               Disable target/bg ratio filter
 --dump                        Dump all the intermediate data, choices: True, False (default: False)
 --output_directory OUTPUT_DIRECTORY
                               Output directory (default: current directory)
 --smooth_size SMOOTH_SIZE
                              Size in bp for the smoothing window (default: internal_window_length/4)
 --gene_annotations_filename GENE_ANNOTATIONS_FILENAME
                              Optional gene annotations file from the UCSC Genome Browser in bed format to map each region to its closes gene
 --gene_ids_to_names_filename GENE_IDS_TO_NAMES_FILENAME
                              Optional mapping file between gene ids to gene names (relevant only if --gene_annotation_filename is used)
 --n_processes N_PROCESSES
                              Specify the number of processes to use. The default is #cores available.
 --version                    Print version and exit*
~~~

### Module 3: haystack_tf_activity_plane

~~~
 *haystack_tf_activity_plane [-h]
                                        [MOTIF_MAPPING_FILENAME]
[--motif_mapping_filename
                                        [--output_directory OUTPUT_DIRECTORY]
                                        [--name NAME] [--plot_all] [--version]
                                        haystack_motifs_output_folder
                                        gene_expression_samples_filename
                                        target_cell_type
 positional arguments:
  haystack_motifs_output_folder
                          A path to a folder created by the haystack_motifs utility
  gene_expression_samples_filename
                          A file containing the list of sample names and locations
  target_cell_type        The sample name to use as a target for the analysis
 optional arguments:
  -h, --help              show this help message and exit
  --motif_mapping_filename MOTIF_MAPPING_FILENAME
                          Custom motif to gene mapping file (the default is for
                          JASPAR CORE 2016 database)
  --output_directory OUTPUT_DIRECTORY
                           Output directory (default: current directory)
  --name NAME              Define a custom output filename for the report
  --plot_all               Disable the filter on the TF activity and correlation (default z-score TF>0 and rho>0.3)
  --version                Print version and exit.*
~~~

## References

Bernstein, B. E., Stamatoyannopoulos, J. A., Costello, J. F., Ren, B., Milosavljevic, A., Meissner, A.,… Thomson, J. A. (2010). The NIH Roadmap Epigenomics Mapping Consortium. Nature Biotechnology, 28(10), 1045–1048.

Dunham, I., Kundaje, A., Aldred, S. F., Collins, P. J., Davis, C. A., Doyle, F.,… Birney, E. (2012). An integrated encyclope dia of DNA elements in the human genome. Nature, 489(7414), 57–74.

Grant, C. E., Bailey, T. L., & Noble, W. S. (2011). FIMO: Scanning for occurrences of a given motif. Bioinformatics, 27(7), 1017–1018.

Jenuwein, T. (2001). Translating the Histone Code. Science, 293(5532), 1074–1080.

Mathelier, A., Fornes, O., Arenillas, D. J., Chen, C. Y., Denay, G., Lee, J.,… Wasserman, W. W. (2016). JASPAR 2016: A major expansion and update of the open-access database of transcription factor binding profiles. Nucleic Acids Research, 44(D1), D110–D115.

Pinello, L., Xu, J., Orkin, S. H., & Yuan, G.-C. (2014). Analysis of chromatin-state plasticity identifies cell-type-specific regulators of H3K27me3 patterns. Proceedings of the National Academy of Sciences, 111(3), E344–E353.

